# A Sequence Labeling Framework for Extracting Drug-Protein Relations from Biomedical Literature

**DOI:** 10.1101/2022.03.31.486574

**Authors:** Ling Luo, Po-Ting Lai, Chih-Hsuan Wei, Zhiyong Lu

## Abstract

Automatic extracting interactions between chemical compound/drug and gene/protein is significantly beneficial to drug discovery, drug repurposing, drug design, and biomedical knowledge graph construction. To promote the development of the relation extraction between drug and protein, the BioCreative VII challenge organized the DrugProt track. This paper describes the approach we developed for this task. In addition to the conventional text classification framework that has been widely used in relation extraction tasks, we propose a sequence labeling framework to drug-protein relation extraction. We first comprehensively compared the cutting-edge biomedical pre-trained language models for both frameworks. Then, we explored several ensemble methods to further improve the final performance. In the evaluation of the challenge, our best submission (i.e., the ensemble of models in two frameworks via major voting) achieved the F1-score of 0.795 on the official test set. Further, we realized the sequence labeling framework is more efficient and achieves better performance than the text classification framework. Finally, our ensemble of the sequence labeling models with majority voting achieves the best F1-score of 0.800 on the test set.

**Database URL:** https://github.com/lingluodlut/BioCreativeVII_DrugProt

## 1 Introduction

A significant number of associations and interactions between chemical/drug and gene/protein have been published in scientific literature. Such unstructured texts, however, are difficult to summarize in a structured format for automatic data analysis. Although a significant amount of manual curation has been involved in summarizing and depositing the information from the literature (e.g., CTD (1) and ChemProt (2)), the process is extremely time-consuming. Given the rapid growth of the literature, developing computational methods to accelerate the efficiency of the knowledge extraction is critical (3, 4). Drugs and proteins play important roles in metabolism and the regulation of biological processes. Extracting the relations between drugs and proteins from the biomedical literature is crucial to various biomedical tasks such as drug discovery, drug repurposing, drug-induced adverse reactions, and biomedical knowledge graph construction. Such extraction is important not only for biological purposes but also to pharmacological and clinical research. In this regard, certain types of relations are highly relevant, including those among metabolic relations, antagonists, agonists, inhibitors, and activators.

Over the past decade, various computational methods have been proposed to extract biomedical relations from text, including early pattern-based methods (5, 6), traditional machine learning methods (7–9), and recently deep learning methods (10–13). For example, Corney et al. developed BioRAT, a template-based information extraction tool, to extract biological information from full-length papers (5). Kim et al. proposed a rich feature-based linear kernel approach to identify drug-drug interactions (DDIs) (9). Zhang et al. proposed a hybrid model that combines recurrent neural network (RNN) and convolutional neural network (CNN) for biomedical relation extraction. Recently, biomedical pre-trained deep learning models (e.g., BioBERT (13)) have achieved the state-of-the-art performance and have become prevalent in the biomedical relation extraction task.

To accelerate the advancement of a method to extract the relations between chemicals and proteins, the BioCreative VI organized the CHEMPROT task (14) for chemical-protein relations. The task requires the determination of whether an interaction exists between a chemical and protein pair in the given text (most are in a sentence), and identifying the granular categories of the relation. Recently, in BioCreative VII, a related task was organized (i.e., BioCreative VII DrugProt (15)), which significantly increased the corpus from 2,432 to 5,000 abstracts. Further, DrugProt increased the number of granular relation categories from 5 to 13 to evaluate the performance of the system.

Most of the participants in BioCreative VI applied deep learning-based neural network methods (e.g., CNNs and RNNs) to extract the relation pairs of chemicals and proteins (14). During the BioCreative VI challenge, the ensemble of support vector machine (SVM) and deep learning models achieved the best performance, with an F1-score of 0.641 (16). Later, transformer-based pre-trained language models, such as BERT (17), have shown the capability of contextual representation at large volumes. Some biomedical versions of the transformer-based model (such as BioBERT (13), PubmedBERT(18), and BioM-ALBERT(19)) were proposed and applied in the biomedical domain. These biomedical pre-trained models show promising results, and these models significantly outperformed the previous state-of-the-art methods. One of the recently published methods, BioM-ALBERT achieves an F1-score of 0.793 on the ChemProt corpus of the BioCreative VI challenge task (19). In general, most of these methods treat the relation extraction task as a text classification problem. A given text with the highlighted chemical and protein is provided to the machine learning-based classifier to recognize the predefined relation types (or no relation). This type of method is required to process all of the pairs between two entities, one by one, which is time-consuming and difficult to handle large-scale data using advanced deep learning techniques. Moreover, these methods ignore the dependency between multiple relations, as they deconstruct the relation extraction into multiple independent relation classification subtasks.

To address these problems, we propose a sequence labeling framework for the drug-protein relation extraction task. In our framework, we convert the task to a sequence labeling problem. Different from the conventional classification framework, our framework can recognize all possible relevant entities (named as tail entity) associated with the given entity (named as head entity) at once. It is more efficient and is able to fully exploit the dependencies of relations for improved performance. We also investigated multiple biomedical pre-trained language models (PLMs) for both frameworks. Further, we applied several ensemble methods to optimize the final performance. The experimental results show that the ensemble of our sequence labeling models with major voting achieves the highest F1-score of 0.800 on the test set.

## 2 Materials and methods

### 2.1 Dataset

In the BioCreative VII DrugProt track, the organizers released a large manually labeled corpus that included annotations of mentions of drugs (including chemical compounds and drugs) as well as proteins (including genes, proteins, and miRNA) and their relations. For benchmarking, the DrugProt corpus was split into training, development, and test sets. In addition, a background set containing 10,000 abstracts with automatic mention annotations was mixed with the test set. During the challenge, the participants are required to process the entire collection (10,000 background abstracts + 750 Gold Standard abstracts) to avoid the potential manual correction of their results. But the final performance was evaluated on the 750 gold standard abstracts only. Further, around 2,400,000 PubMed records with automatic mention annotations were provided to the participants in the additional DrugProt Large Scale subtask (15). Table 1 shows an overview of the DrugProt corpus.

**Table 1.**
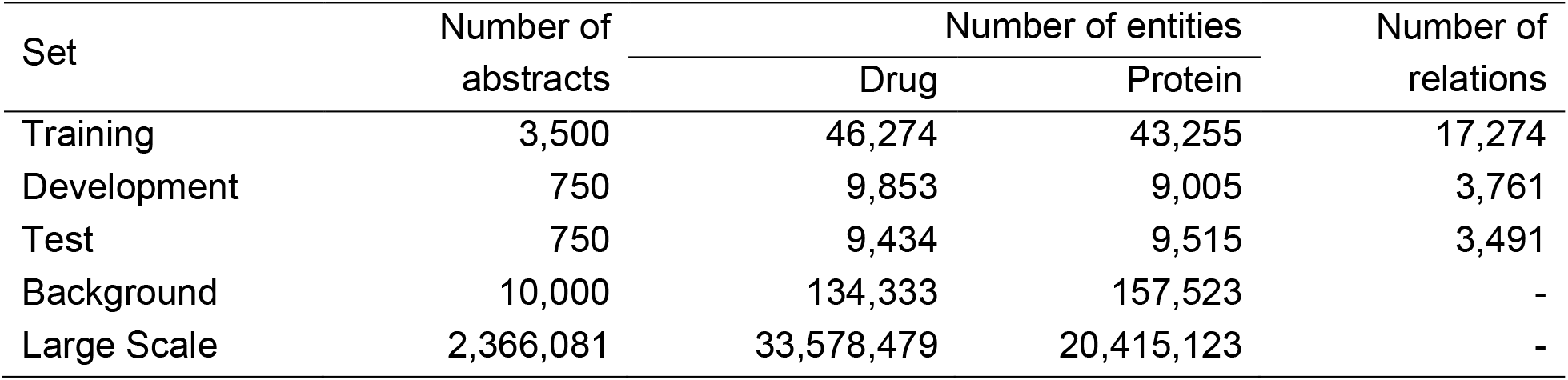
The overview of DrugProt corpus

The DrugProt corpus defined 13 relation types, including INDIRECT-DOWNREGULATOR, INDIRECT-UPREGULATOR, DIRECT-REGULATOR, ACTIVATOR, INHIBITOR, AGONIST, ANTAGONIST, AGONIST-ACTIVATOR, AGONIST-INHIBITOR, PRODUCT-OF, SUBSTRATE, SUBSTRATE_PRODUCT-OF, and PART-OF. Figure 1 shows the population of the relation types in the training and development merged set. The distribution of the relation types is highly imbalance. The most frequent relation type is INHIBITOR with over 6,000 instances, but some other relation types have fewer than 50 instances (e.g., AGNONIST-INHIBITOR). As the characteristic we observed in the corpus, cross-sentence relations are rare, appearing in less than 1% of the training set. Therefore, our developed method focuses on the relation extraction challenge within a single sentence. For the cross-sentence relations in the training set, we merged the continued sentences describing the relation into one sentence.

**Figure 1.**
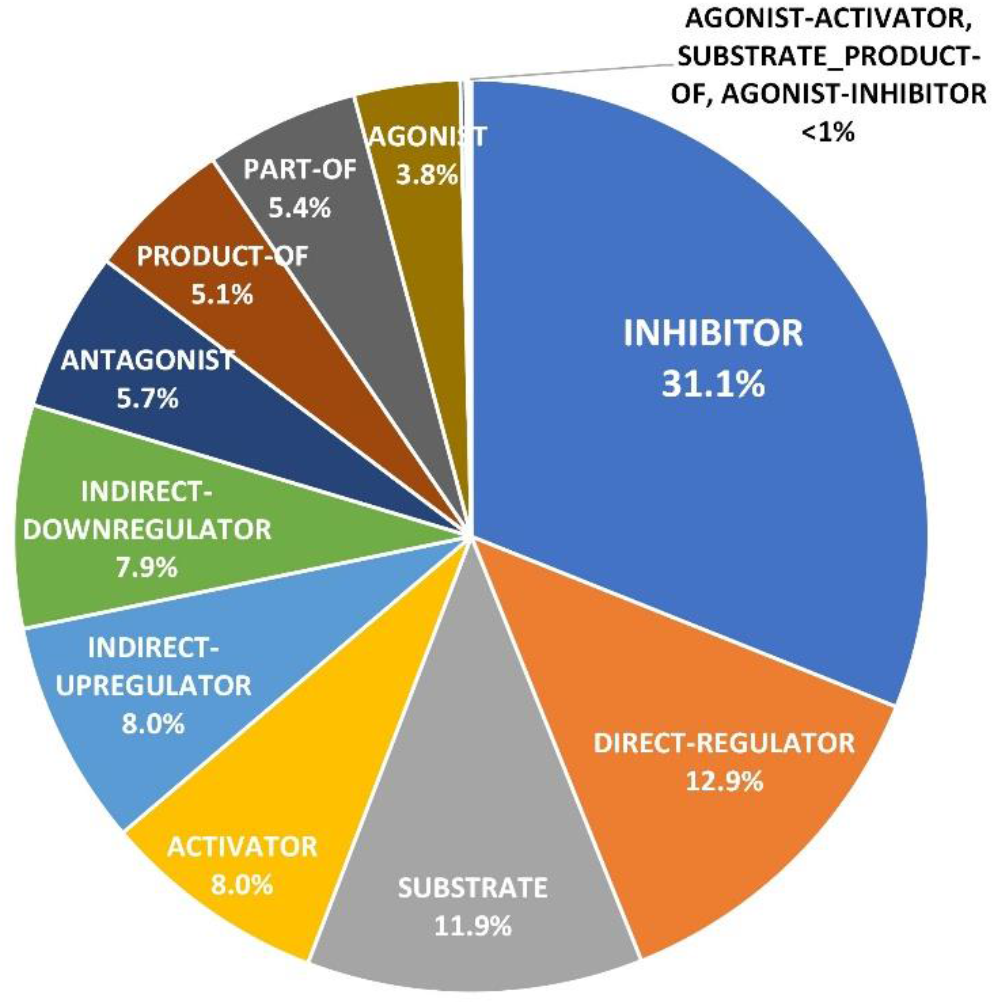
The distribution of the relation categories in the DrugProt in the training and development merged set.

### 2.2 Text classification framework

To provide a comparison with our proposed sequence labeling framework, we also built the corresponding neural network models in a conventional text classification framework with minimal architectural modification. The two frameworks are illustrated in Figure 2. We first introduce the basic text classification framework for this task as part of our methods. In this framework, the relation extraction task is treated as a multi-class classification problem. As shown in Figure 2A, we first need to generate all drug-protein entity pairs in the sentence as the input instances and then process all entity pairs one by one. In each input instance, we inserted the “<Argl></Argl>” and “<Arg2></Arg2>” tag pairs to the front and end of the head- and tail-entity, the “<Drug></Drug>” tag pairs to the other drugs, and the “<Prot></Prot>” tag pairs to the other proteins. Given the input instance with the candidate entity pair, we built a classifier to classify the entity pair into a predefined relation type (or no relation). We used the biomedical pre-trained language model (PLM) to encode the input text. Then the [CLS]’s output vector at the last hidden layer of the PLM is fed into a fully connected layer with ReLU (20) activation function. Finally, a softmax classification layer is used to classify the relation of an entity pair. In the experiments, we evaluated the five biomedical PLMs including PubMedBERT (18), BioBERT (13), BioRoBERTa (21), BioM-ELECTRA (19), and BioM-ALBERT (19) to this task.

**Figure 2.**
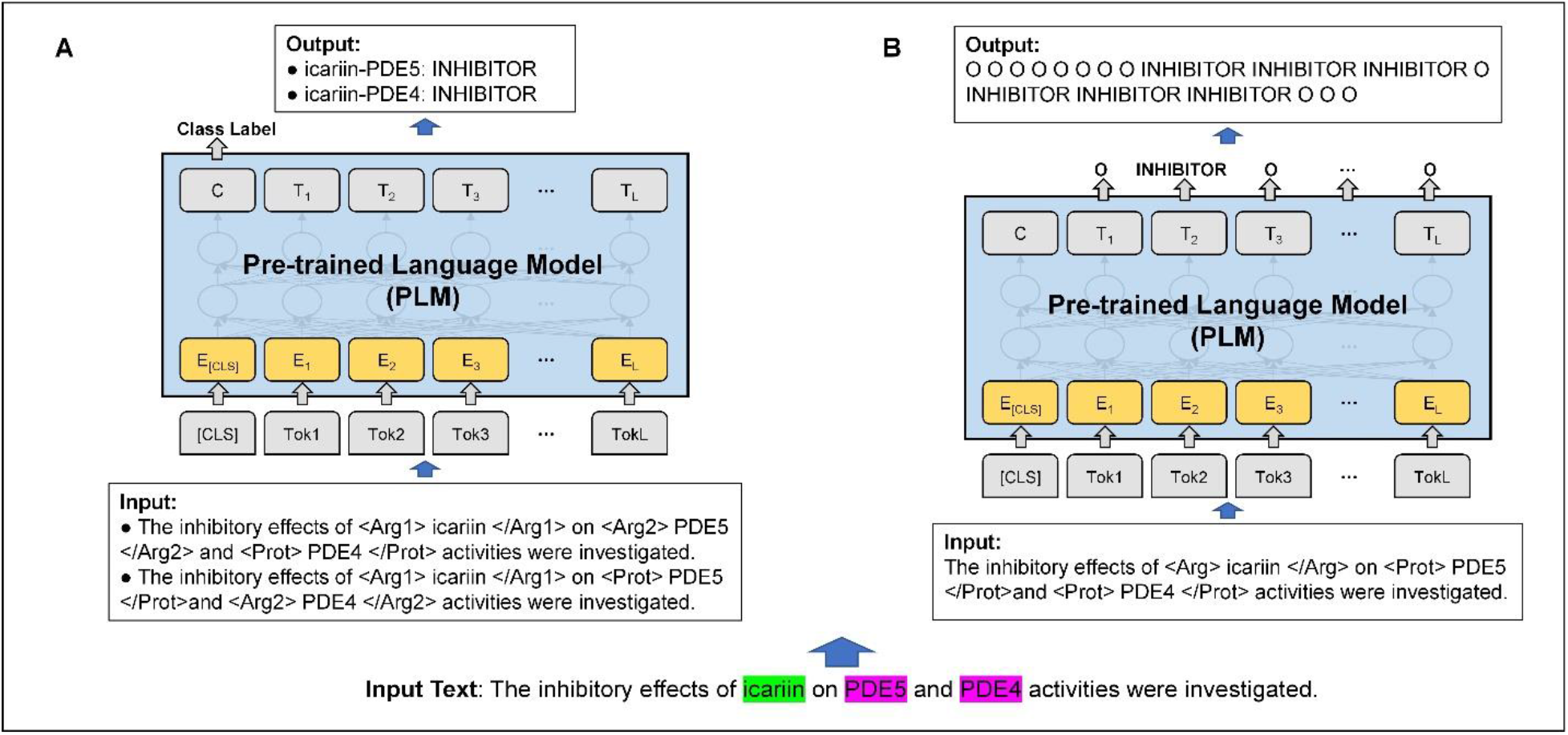
Example of relation extraction in text classification and sequence labeling frameworks. **(A)** Text classification framework. **(B)** Sequence labeling framework. In the example, the drug of “icariin” has INHIBITOR relations with the proteins of “PDE5” and “PDE4”.

### 2.3 Sequence labeling framework

Inspired by our previous work (22, 23), we proposed a sequence labeling framework to address the sentence-level biomedical relation extraction task. As shown in Figure 2B, we converted the task to a sequence labeling problem. Given a head entity (e.g., drug entity of “icariin”) in the sentence, the goal of the model is to recognize all the corresponding tail entities (e.g., protein entities of “PDE5” and “PDE4”) that are involved in the drug-protein relations with the head entity. We defined two different labeling strategies from different perspectives to extract the relation between drug and protein, including (1) from drug to protein (D→P), the strategy extracts the protein tail entities by the given drug head entity; and (2) from protein to drug (P→D), the strategy extracts the drug tail entities by the given protein head entity. As an example shown in Figure 2, the text classification framework deconstructed input text with two entity pairs into two independent sentence classification subtasks, while our sequence labeling framework can effectively formulate the problem to one sequence labeling task. Therefore, our sequence labeling framework is able to fully exploit the dependencies of the given head entity and all corresponding tail entities in one instance, which is more efficient. Next, we use D→P strategy as the example to describe the details of our method.

#### (1) Tagging scheme

We designed a simple and effective tagging scheme for this task. We firstly refined the input sequence by inserting the “<Arg></Arg>” tag pair to locate the boundary of the drug head entity, the “<Drug></Drug>” tag pairs to the other drugs, and “<Prot></Prot>” tag pairs to the proteins. For the output sequence, each token is assigned a label that contributes to the extraction. The tokens involved in the relations are tagged by the corresponding relation type labels (e.g., INHIBITOR), which are predefined according to the training sets. We defined the label “O” to represent other tokens that do not involve in a relation. Figure 2B shows an example of tagging a sentence with our tagging scheme, according to the original gold standard annotations of the DrugProt dataset. In this example, the input sentence contains three entities (i.e., the drug entity of “icariin” and the protein entities of “PDE5” and “PDE4”) and two drug-protein relation triples (i.e., {icariin, INHIBITOR, PDE5} and {icariin, INHIBITOR, PDE4}). We selected the drug of “icariin” as the head entity to predict the corresponding protein entities associated with the drug. We inserted the “<Arg>” and “</Arg>” tags in the front and end of the head entity “icariin” and added the entity type tags of “<Prot></Prot>” to the protein entities of “PDE5” and “PDE4”. Because the “PDE5” and “PDE4” participate in the relation of “INHIBITOR” with the head entity “icariin”, the output labels of those proteins are “INHIBITOR”, and other tokens are “O”.

#### (2) Model training

Based on our tagging scheme, we first transformed the training data into the training instances. Then we selected several cutting-edge PLMs to fine-tune for this task. Recently, transformer-based pre-trained models have shown promising results in a broad range of natural language processing (NLP) tasks and are widely used in the sequence labeling tasks (such as named entity recognition) (24). A large array of pre-trained models that are pre-trained on PubMed abstracts and PMC full-text articles are available in the biomedical domain. With minimal architectural modification, biomedical PLMs can be applied to various downstream biomedical text mining tasks and can significantly outperform previous state-of-the-art models in terms of biomedical NLP tasks (25). Given an input instance, we used biomedical PLMs as the encoder to represent the text. To optimize the performance, we added a fully connected layer with the ReLU activation function to finalize the hidden representation of the biomedical PLM for each token. Then, we used a softmax classification layer over the output label set to predict the label probability score of each token. In the experiments, we evaluated the same five biomedical PLMs (i.e., PubMedBERT, BioBERT, BioRoBERTa, BioM-ELECTRA, and BioM-ALBERT) in the text classification framework to the sequence labeling method. The training objective is to minimize the cross-entropy loss. In our sequence labeling framework, the number of tokens with the “O” class label is much large than the number of tokens with relation labels since there are always many more non-relation tokens in a sentence. Therefore, in addition to the standard categorical cross-entropy loss, we further explored sample weights in the loss function to deal with the class imbalance issue. Here the samples for class *C* are weighted by the equation: *W_c_* = log (total number of the samples/number of samples in class *C*). Our experimental results, however, show that the weighted loss does not achieve better performance than the standard loss.

#### (3) Model prediction

In the prediction phase, the input text was first split into sentences and tokenized using the Stanza tool (26). Only the sentences with both drug and protein entities were used to extract the relations. In each iteration, we sequentially selected a drug entity as the current head entity and assigned the tag pairs by the tagging scheme to distinguish the current drug from the other drug and protein entities. Then we use the trained model to tag the tokens of the sentence to extract the corresponding proteins associated with the given drug. If a conflict of the relation types exists between the tokens of the tail entity, the label of the first token of the tail entity would be selected as the final relation type.

### 2.4 Model ensemble

To further optimize the performance, three ensemble alternatives (i.e., majority voting, voting with random search, and voting with backward search) were investigated to combine the results of the different models in our experiments. For majority voting, we selected the relation class that is predicted by more than half of all models. In addition, we searched backward and randomly to find a subset of our models that might achieve higher performance on the development set, rather than using all models. In the random search, we randomly generated a combination of our models each time, and we kept the best performance on the development set until the number of combinations reached our predefined value. In the backward search, we combined the results of all models. Then we iteratively removed the models to achieve higher performance until we found the combination of the models with the highest performance on the development set.

## 3 Results and discussion

### 3.1 Experimental setting

We downloaded five biomedical PLMs (i.e., PubMedBERT^1^, BioBERT^2^, BioRoBERTa^3^, BioM-ELECTRA^4^, and BioM-ALBERT^5^) and evaluated them in both frameworks. Note that, with the exception of PubMedBERT, which has only a base version, the other PLMs have the base and large versions. The large PLMs were used as our final selection, as they achieved better performance than the base models in our experiments. For the parameter setting, we used PLMs with the default parameter settings and set the main hyper-parameters as follows: learning rate of 1e-5, batch size of 16, and last fully connected layer size of 128. The number of training epochs was chosen by the early stopping strategy (27) (50 epochs at most). Specially, we explored three methods: (1) DevES, whereby the training set is used for model training, and the development set is used for early stopping according to the overall F1-score; (2) ValES, for which we merged the official training and development sets, then randomly selected 350 abstracts as our validation set for early stopping according to the overall F1-score, and the remaining abstracts were used as the training set; and (3) TrainES, for which we merged the official training and development sets for model training, then chosen number of training epochs by early stopping according to the training loss score. Our models are implemented using the open-source deep learning libraries: HuggingFace’s Transformer (28) and TensorFlow (29). The performance of the models is evaluated on the test set using the official evaluation script with micro-averaged scores: precision (P), recall (R), and F1-score (F1).

### 3.2 Performance of individual models in sequence labeling framework

Table 2 presents the results of individual models with different PLMs and different labeling strategies in the sequence labeling framework on the test set. All models were trained using the standard categorical cross-entropy loss with the DevES way for fair comparisons.

**Table 2.**
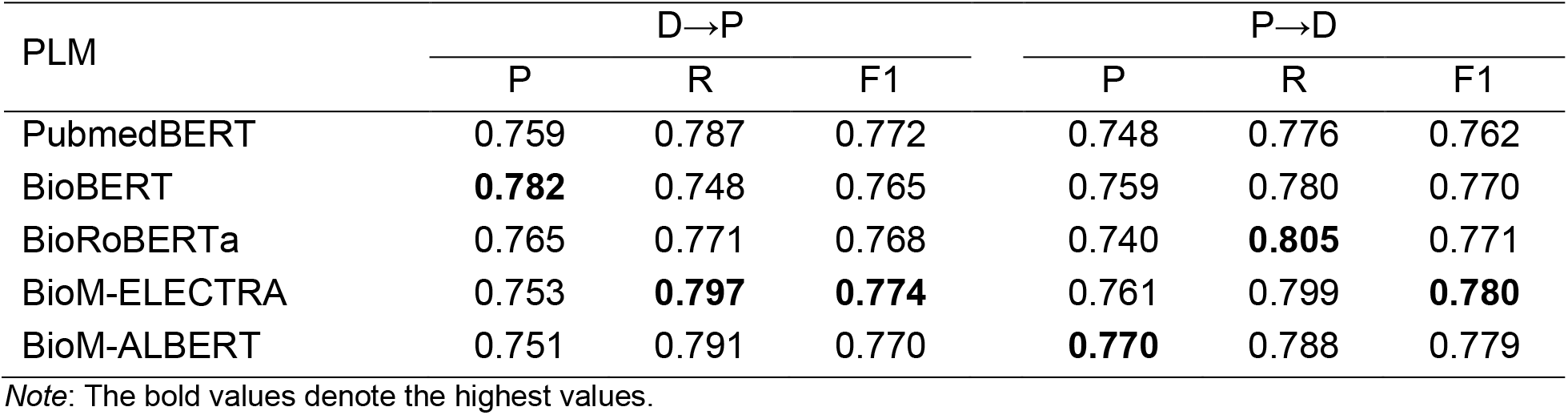
Performance of individual models in sequence labeling framework on the test set

| PLM | D→P |  |  | P→D |  |  |
| --- | --- | --- | --- | --- | --- | --- |
|  | P | R | F1 | P | R | F1 |
| PubmedBERT | 0.759 | 0.787 | 0.772 | 0.748 | 0.776 | 0.762 |
| BioBERT | **0.782** | 0.748 | 0.765 | 0.759 | 0.780 | 0.770 |
| BioRoBERTa | 0.765 | 0.771 | 0.768 | 0.740 | **0.805** | 0.771 |
| BioM-ELECTRA | 0.753 | **0.797** | **0.774** | 0.761 | 0.799 | **0.780** |
| BioM-ALBERT | 0.751 | 0.791 | 0.770 | **0.770** | 0.788 | 0.779 |

*Note:* The bold values denote the highest values.

A comparison of the two sequence labeling strategies (D→P and P→D) shows that most PLMs with P→D achieve better F1-scores than the same PLMs with D→P, except PubmedBERT. The average F1-score of P→D is slightly higher than that of D→P (0.772 vs. 0.770). Further, in a comparison of the different PLMs in the same sequence labeling strategy, BioM-ELECTRA achieves the highest F1-scores using both strategies (0.774 and 0.780 of D→P and P→D, respectively). Other models obtained similar performance.

### 3.3 Comparison of sequence labeling and text classification frameworks

To explore the effectiveness of our sequence labeling framework, we comprehensively compared the performance of individual models in classification and sequence labeling frameworks on the test. All models are trained using the standard categorical cross-entropy loss by DevES method. We selected the results of the model with a better F1-score in two sequence labeling strategies as the sequence labeling result. The performance comparison is shown in Figure 3.

**Figure 3.**
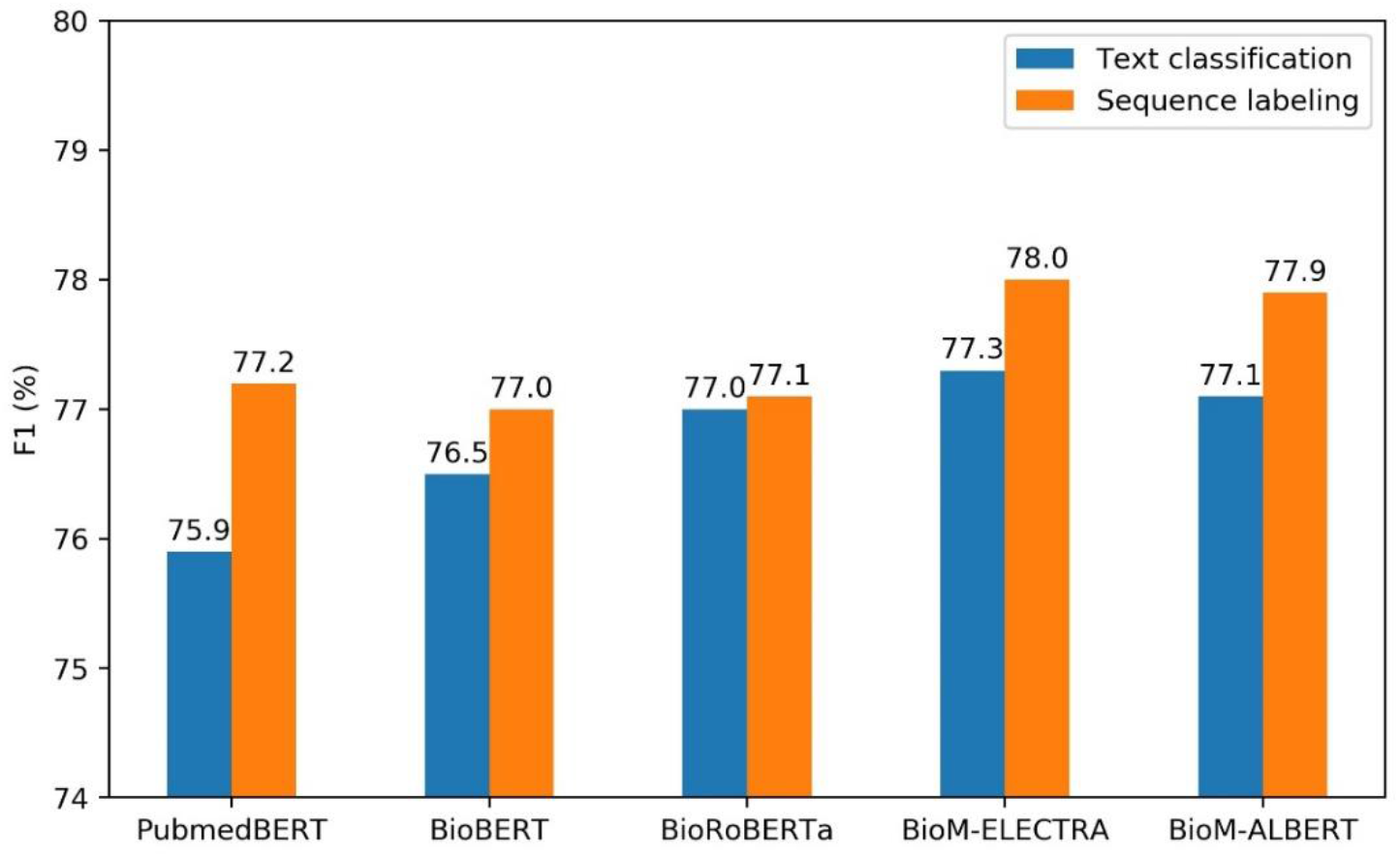
Performance comparison of the text classification and sequence labeling frameworks

The results of the models in the text classification framework show a similar trend to that of the models in the sequence labeling framework. BioM-ELECTRA achieves a slightly better performance than the other PLMs in the text classification framework, with an F1-score of 0.773. Compared with the text classification framework, all models in our sequence labeling framework achieve better performance (an average improvement of 0.68% in F1-score). Compared with the text classification framework, the sequence labeling framework can learn more dependency information between the head entity and all tail entities. The text classification framework ignores the dependency, as all entity pairs are classified independently.

Moreover, we recorded the prediction processing time of the PLMs in the different frameworks per 100 abstracts to compare the processing times of the two frameworks as shown in Table 3. All models are tested on the same GPU (Tesla V100-SXM2-32GB) and CPU (Intel(R) Xeon(R) Gold 6226 CPU @ 2.70GHz, 24Cores). The results show that our sequence labeling framework is more efficient than the text classification framework on both GPU and CPU. The main reason is that the model in the text classification framework is required to process all the pairs between drug and protein entities one by one, but the sequence labeling framework can recognize all possible tail entities associate with the given head entity at once. For example, for a sentence with *N* drug entities and *M* protein entities, the relation extraction task is deconstructed into *N* × *M* sentence classification subtasks in a conventional classification framework. Our sequence labeling (D→P) can effectively narrow this down to *N* sequence labeling subtasks, and our sequence labeling (P→D) can narrow this down to M sequence labeling subtasks. As shown in Table 1, the numbers of drug and protein entities are approximate in the DrugProt corpus. Therefore, the time complexity dropped from O(*N* × *M*) to O(*N*) or O(*M*). As we observed, the processing time of the two sequence labeling strategies is similar, and it is ~2 times faster than the text classification framework. This indicates that selecting the type of entities with a lower number of entities as the head entity is more efficient in our sequence labeling framework.

**Table 3.**
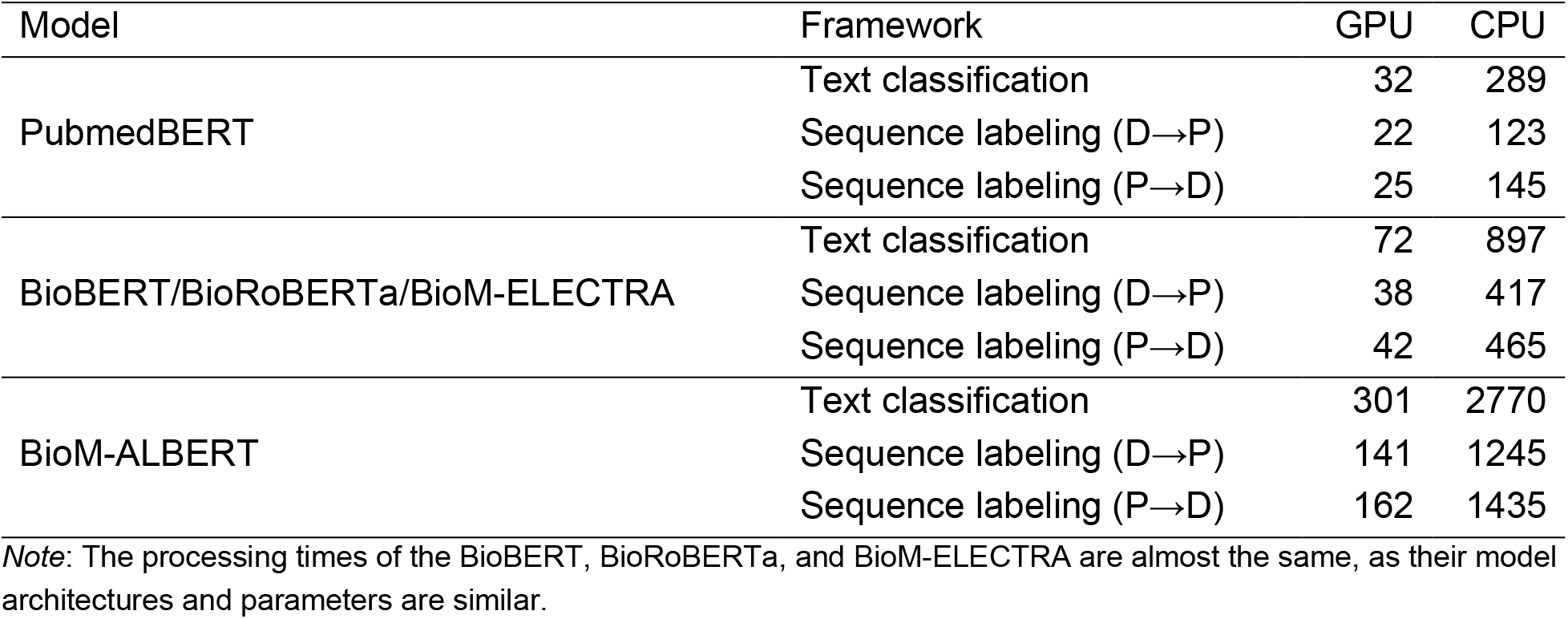
Processing time comparison (Seconds per 100 abstracts)

### 3.4 Performance of different model combinations

To further improve performance, we explored three alternatives (i.e., majority voting, voting with random search, and voting with backward search), as described in the “Model Ensemble” section, to combine the individual model results. During the DrugProt challenge, we submitted the following five runs as our final submissions:

- Run 1: The ensemble of all sequence labeling models, including four types (the combinations of the D→P and P→D with standard loss and weighted loss) for each PLM, trained by ValES via majority voting.
- Run 2: The ensemble of all sequence labeling models trained by DevES via majority voting.
- Run 3: The ensemble of the sequence labeling and text classification models trained by DevES via voting with backward search.
- Run 4: The ensemble of the sequence labeling and text classification models trained by DevES via voting with random search.
- Run 5: The ensemble of all sequence labeling and text classification models trained by DevES and ValES via majority voting.

Run 5 (i.e., the ensemble of all models) achieves slightly better performance and obtains the highest overall F1 -score on the test set in our five submissions. Comparing the different ensemble methods, although the voting with random search and backward search ensemble method achieves better performance on the development set, they do not achieve better performance than the simple major voting on the test set. We found that, compared with DevES, the models with the early stopping strategy of ValES achieve worse performance. One possible reason is that the size of our validation set (350 abstracts) is small so that the performance of the models trained with the ValES is unstable. Therefore, at post-challenge, we trained the models with TrainES instead of ValES to make full use of the development set. Moreover, to investigate the effect of sample weighting in the loss function, we compared the sequence labeling models with standard loss and weighted loss. Thus, we recombined the models with majority voting based on the following configurations:

- Run 6: The ensemble of all text classification models trained by DevES via majority voting.
- Run 7: The ensemble of all sequence labeling models with standard loss trained by DevES via majority voting.
- Run 8: The ensemble of all sequence labeling models with weighted loss trained by DevES via majority voting.
- Run 9: The ensemble of all text classification models trained by DevES and TrainES via majority voting.
- Run 10: The ensemble of all sequence labeling models with standard loss trained by DevES and TrainES via majority voting.
- Run 11: The ensemble of all text classification and sequence labeling models with standard loss trained by DevES and TrainES via majority voting.

From the results shown in Table 4, as compared with the best results of individual models, it is clear that the ensemble of models with majority voting (i.e., Runs 6 and 7) improves performance (corresponding improvements of 1.5% and 1.0% in sequence labeling and text classification frameworks, respectively). The main reason is that different kinds of PLMs may contribute diverse information from different perspectives and an ensemble of them can improve the performance. Moreover, a single PLM sometime is unstable due to some randomness (such as random initialization of parameters). Ensembling multiple models can improve the robustness and reliability in the performance. Comparing Run 7 and 8, we observed that the model trained using the weighted loss function achieves better recall but worse precision than does the model trained using the standard loss function. The results show the model with a weighted loss function can predict more relation labels but also bring some wrong relation labels leading thus it cannot obtain a higher F1-score than the model with the standard loss. Ensemble of the modes with weighted loss and standard loss also cannot obtain better performance. In future work, we will further explore the label imbalance problem. When the models trained with TrainES are added into the combination, the performances are further improved. Compared with the ensemble of text classification models, the ensemble of sequence label models achieves better performance. Finally, we found that Run 10 (i.e., the ensemble of only sequence labeling models trained by DevES and TrainES with standard loss) achieves the best performances in all runs, even better than the ensemble of all models (Run 11).

**Table 4.** Performance of different model combinations on the test set

| Method | Framework | Early stopping | Ensemble | P | R | F1 |
| --- | --- | --- | --- | --- | --- | --- |
| Single | BioM-ELECTRA in TC | DevES | - | 0.774 | 0.773 | 0.773 |
| Single | BioM-ELECTRA in SL (standard loss) | DevES | - | 0.761 | 0.799 | 0.780 |
| Run 1 | SL (standard and weighted loss) | ValES | Majority voting | 0.782 | 0.799 | 0.791 |
| Run 2 | SL (standard and weighted loss) | DevES | Majority voting | 0.793 | 0.795 | 0.794 |
| Run 3 | SL (standard and weighted loss) + TC | DevES | Backward search | 0.785 | 0.803 | 0.794 |
| Run 4 | SL (standard and weighted loss) + TC | DevES | Random search | 0.790 | 0.798 | 0.794 |
| Run 5 | SL (standard and weighted loss) + TC | DevES + ValES | Majority voting | 0.785 | **0.805** | 0.795 |
| Run 6 | TC | DevES | Majority voting | 0.784 | 0.782 | 0.783 |
| Run 7 | SL (standard loss) | DevES | Majority voting | 0.809 | 0.781 | 0.795 |
| Run 8 | SL (weighted loss) | DevES | Majority voting | 0.786 | 0.799 | 0.792 |
| Run 9 | TC | DevES + TrainES | Majority voting | 0.801 | 0.775 | 0.788 |
| Run 10 | SL (standard loss) | DevES + TrainES | Majority voting | **0.802** | 0.797 | **0.800** |
| Run 11 | SL (standard loss) + TC | DevES + TrainES | Majority voting | 0.801 | 0.793 | 0.797 |

*Note:* SL denotes sequence labeling framework; TC denotes text classification framework. The bold values denote the highest values.

### 3.5 Performance comparison with other existing methods

To further demonstrate the effectiveness of our approach, we compare our methods with the other two teams of the top three in the BioCreative VII DrugProt challenge. Table 5 shows the detailed granular results by relation type (F1-score for each relation type) and the overall results of the methods on the test set. Among these methods, both Humboldt (30) and KU-AZ (31) teams’ methods are the ensemble of PLMs based on the text classification framework. The result of NLM-NCBI (32) is the result of our best submission during the challenge (i.e., Run 5). NLM-NCBI-single is the best result of our single model (i.e., the BioM-ELECTRA (P→D) in the sequence labeling framework), and NLM-NCBI-ensemble is our best result of the ensemble (i.e., Run 10, the ensemble of only sequence labeling models with standard loss trained by DevES and TrainES) which was finalized after the challenge.

**Table 5.**
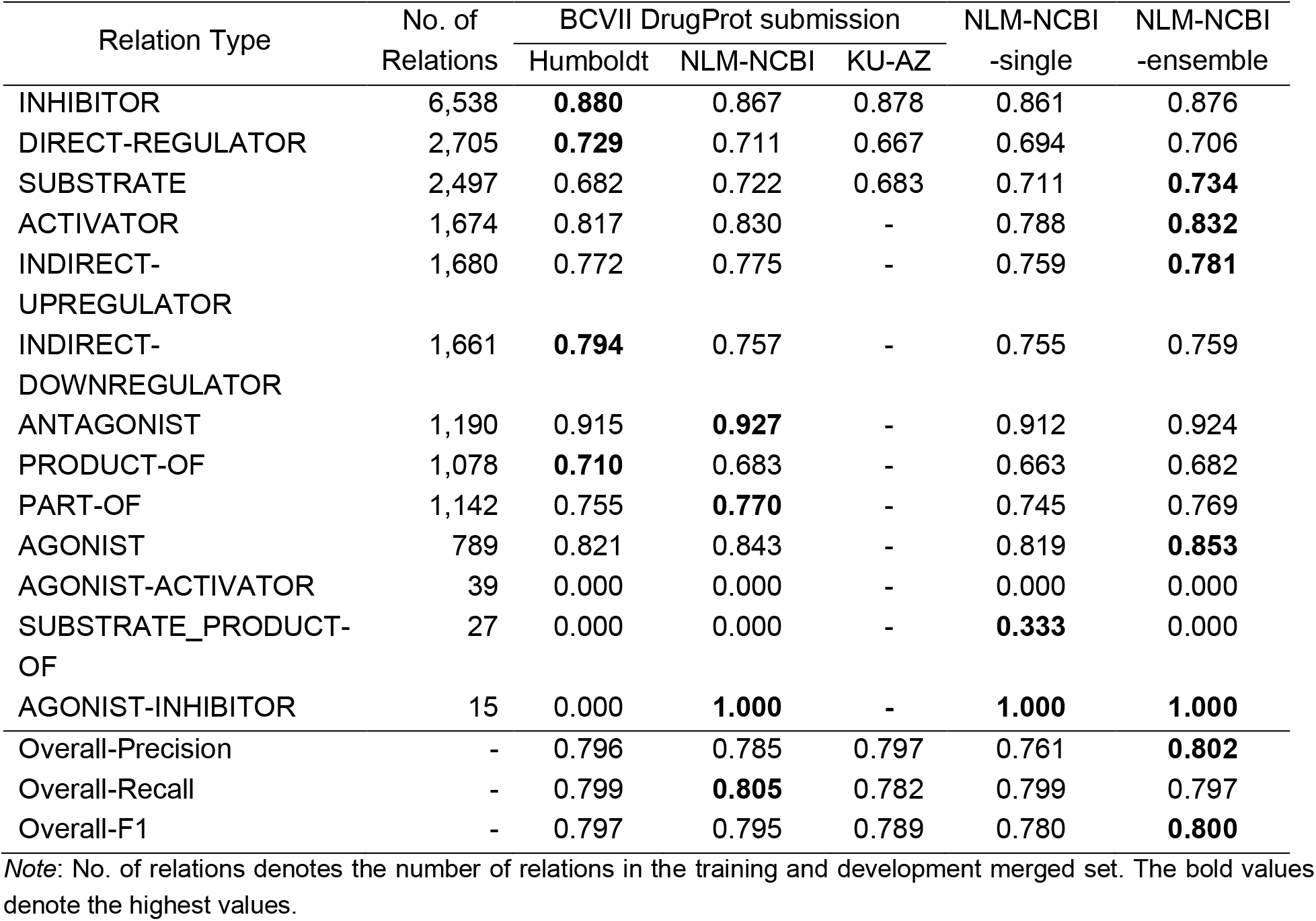
Detailed granular results on the test set

Both the Humboldt and KU-AZ teams explored the effect of the addition database (i.e., Comparative Toxicogenomics Database, CTD (1)) for this task. The Humboldt team used the chemical definitions in the CTD chemical vocabulary to enrich the model input. Their experimental results show that the chemical descriptions can cause an improvement of 0.79% in the F1-score (30). The KU-AZ team selected the chemical-gene interactions dump from CTD to refine data augmentation. However, the overall performance of models pre-trained on augmented datasets is not better than the performance of models without the augmented dataset (31). Compared with the results of the single model, all ensemble methods are able to improve performance. Among these methods, our best ensemble of sequence labeling models achieves slightly better performance than the result of the top 1 team. Our method, without any additional datasets or post-processing, achieves an overall F1-score of 0.800. The detailed granular results by relation type show that the F1-score of our method on the ANTAGONIST relation is higher than 0.900, and F1-scores of most relation types with enough training data are over 0.750. Notably, relation types such as AGONIST-ACTIVATOR and SUBSTRATE_PRODUCT-OF seem to be more difficult to detect due to the small size of training data. Although PRODUCT-OF, DIRECT-REGULATOR, and SUBSTRATE have large training data support, they have relatively lower metrics. These relation types seem to be more complicated to be detected.

Moreover, we participated in the additional DrugProt large-scale subtask, specifically focusing on the scalability and processing of large datasets. During the challenge, we did not use all models to predict the results, as some large PLMs are computationally expensive on the large-scale test set. Instead, we selected four efficient models (i.e., three sequence labeling models: PubMedBERT, BioM-ELECTRA, and BioRoBERTa; and a text classification model: PubMedBERT), and then used different combinations of them with simple majority voting to generate our submissions. Each model took ~5 days to predict the whole large-scale test set (2,366,081 PubMed abstracts) on one GPU. Finally, our best submission (i.e., the ensemble result of all models other than the PubMedBERT sequence labeling model) achieves the highest F1-score, 0.789, among all submissions.

### 3.6 Error analysis

Although our sequence labeling method exhibits promising results for the DrugProt task, there is still room for improvement. We made statistics for the error cases in which our ensemble of sequence labeling models with major voting on the development set, as the test gold standard annotation was not released. There are 604 false positives and 802 false negatives. Among these, there are 92 cases that correctly identify the association of the entity pair but assign the wrong relation type. More errors are caused by the fact that the gold standard relations cannot be identified from the text. To further understand the possible causes of errors, we randomly sample 100 abstracts from the development set and manually analyzed the error cases. We grouped the errors into several major categories, as shown in Table 6.

**Table 6.**
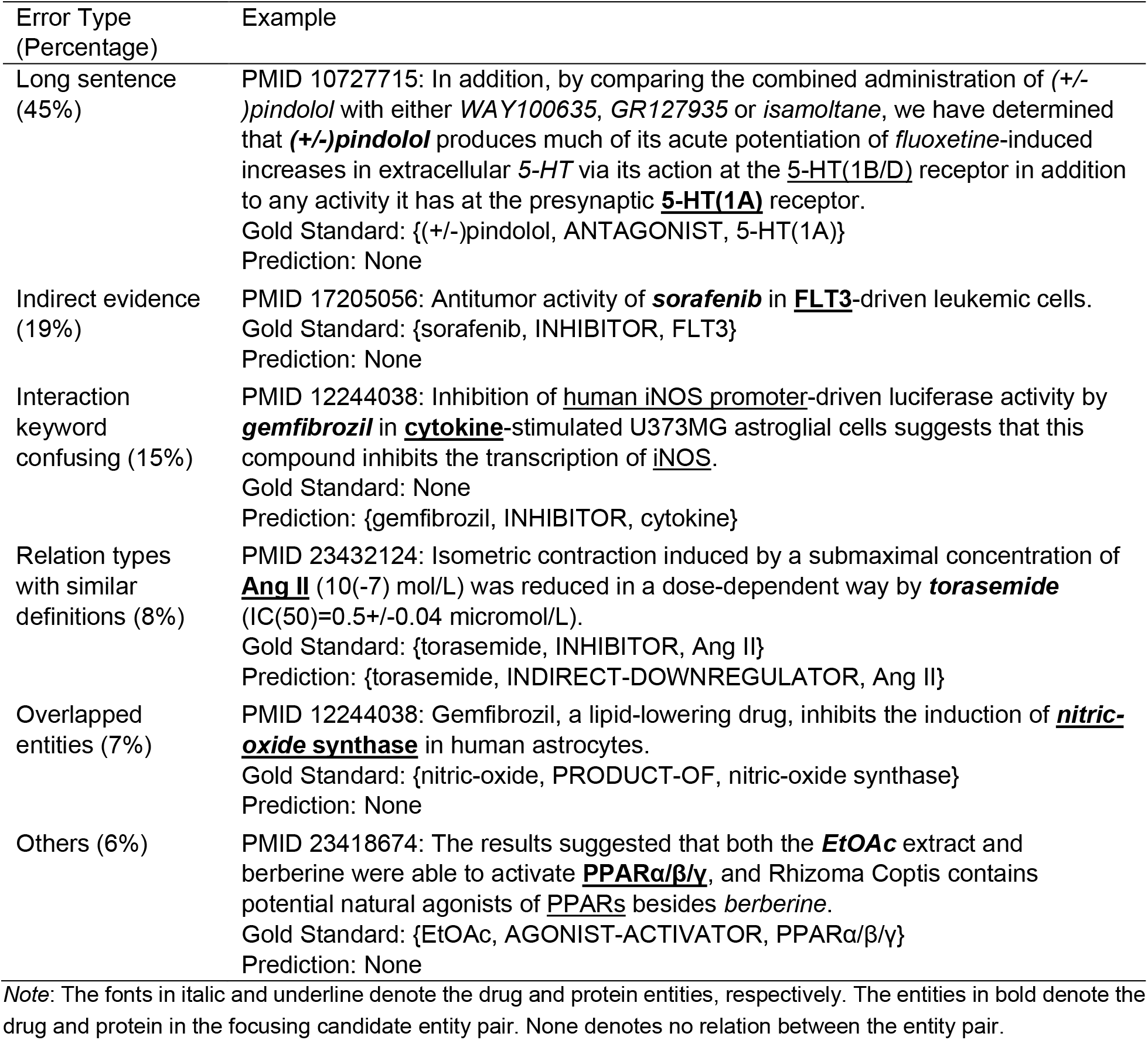
The summary of relation extraction error types

| Error Type<br>(Percentage) | Example |
| --- | --- |
| Long sentence<br>(45%) | PMID 10727715: In addition, by comparing the combined administration of (+/-) *pindolol* with either *WAY100635*, *GR127935* or *isamoltane*, we have determined that (+/-) *pindolol* produces much of its acute potentiation of *fluoxetine*-induced increases in extracellular 5-HT via its action at the 5-HT(1B/D) receptor in addition to any activity it has at the presynaptic 5-HT(1A) receptor.<br>Gold Standard: {(+/-)pindolol, ANTAGONIST, 5-HT(1A)}<br>Prediction: None |
| Indirect evidence<br>(19%) | PMID 17205056: Antitumor activity of *sorafenib* in *FLT3*-driven leukemic cells.<br>Gold Standard: {sorafenib, INHIBITOR, FLT3}<br>Prediction: None |
| Interaction<br>keyword<br>confusing (15%) | PMID 12244038: Inhibition of *human iNOS promoter*-driven luciferase activity by *gemfibrozil* in *cytokine*-stimulated U373MG astroglial cells suggests that this compound inhibits the transcription of *iNOS*.<br>Gold Standard: None<br>Prediction: {gemfibrozil, INHIBITOR, cytokine} |
| Relation types<br>with similar<br>definitions (8%) | PMID 23432124: Isometric contraction induced by a submaximal concentration of *Ang II* (10(-7) mol/L) was reduced in a dose-dependent way by *torasemide* (IC(50)=0.5+/-0.04 micromol/L).<br>Gold Standard: {torasemide, INHIBITOR, Ang II}<br>Prediction: {torasemide, INDIRECT-DOWNREGULATOR, Ang II} |
| Overlapped<br>entities (7%) | PMID 12244038: Gemfibrozil, a lipid-lowering drug, inhibits the induction of *nitric-oxide synthase* in human astrocytes.<br>Gold Standard: {nitric-oxide, PRODUCT-OF, nitric-oxide synthase}<br>Prediction: None |
| Others (6%) | PMID 23418674: The results suggested that both the *EtOAc* extract and berberine were able to activate *PPARα/β/γ*, and *Rhizoma Coptis* contains potential natural agonists of *PPARs* besides *berberine*.<br>Gold Standard: {EtOAc, AGONIST-ACTIVATOR, PPARα/β/γ}<br>Prediction: None |

*Note:* The fonts in italic and underline denote the drug and protein entities, respectively. The entities in bold denote the drug and protein in the focusing candidate entity pair. None denotes no relation between the entity pair.

In our observation, the sentences with longer length bring higher difficulty to the relation extraction task, which caused 45% of the errors. As an example in PMID 10727715, there are 30 words between the drug of “(+/-)pindolol” and the protein of “5-HT(1A)”. Our method missed the ANTAGONIST relation between the two entities, as the long dependency information is difficult to be captured. Another example in PMID 9990013, “This direct biochemical evidence of cooperative interaction in nucleotide binding of the two NBFs of SUR1 suggests that glibenclamide both blocks this cooperative binding of ATP and MgADP and, in cooperation with the MgADP bound at NBF2, causes ATP to be released from NBF1”, contains seven relations among six drug entities and four protein entities, but our method missed four of them and mistakenly extract two relations. As shown in the examples, longer sentences are often with multiple conjunctions or entities, which aggravate the difficulty of the relation extraction. To better improve the performance on those long sentences, the straightforward idea is to incorporate the deep semantics and syntax of the sentence using dependency and constituency parsing which will be considered in our future work. The errors of the second category are caused by the lack of explicit text evidence. Such an example in PMID 17205056, an INHIBITOR relation of the entity pair was manually curated due to the implicit evidence of “Antitumor activity”, but our method cannot recognize it. When evidence is very limited, the additional knowledge base (e.g., CTD) may improve the performance. The third category of the errors is the false positive which incorrectly identifies a relation between two unrelated entities. The model is frequently confused when some frequent keywords (e.g., inhibition) indicating the relations appear nearby the unrelated entities. In PMID 12244038, no relation exists between “gemfibrozil” and “cytokine”, but our model mistakenly recognized the pair as an INHIBITOR relation because the keyword “Inhibition” co-occurs in the sentence. The errors of the fourth category are due to the similar definitions of two relation types (e.g., INHIBITOR and INDIRECT-DOWNREGULATOR), and our method is sometimes confused. Next, the relations between two entities overlapped with each other are harder than the regular entity pairs, especially our sequence labeling framework cannot deal with such entity pairs. The number of remaining errors is relatively small and caused by various reasons, such as the huge gap in the number of instances among different relation types. Some relation types (e.g., AGONIST-ACTIVATOR) are extremely small in number, leading to the model cannot learn enough information to extract these relations correctly. Besides, we also observed a few incorrect annotations in the gold standard dataset.

## 4 Conclusions

In this paper, we present our method, which is based on the pre-trained language models to deal with the challenge of the BioCreative VI DrugProt task. In addition to the classical text classification framework, we propose a sequence labeling framework to extract the relations of drugs and proteins. The experimental results show that our proposed framework is more efficient and is able to fully exploit the dependencies of relations for improved performance. Moreover, the different model combinations by different ensemble methods are further explored to optimize the final performance. The results show that the ensemble of only sequence labeling models with major voting achieves the best performance.

Our sequence labeling framework without any additional knowledge bases or post-processing exhibits promising results for the DrugProt task. In future work, we will investigate whether external resources (e.g., existing knowledge bases, dependency parser information) can be used to further improve our method. In addition, our sequence labeling framework can be easily adapted to other biomedical entity relation extractions. We will test the generalizability of our framework to other biomedical relations such as drug-drug interactions.

## Acknowledgements

The authors thank the organizers of the BioCreative VII DrugProt task. This research was supported by the Intramural Research Program of the National Library of Medicine (NLM), National Institutes of Health.

## Footnotes

1 https://huggingface.co/microsoft/BiomedNLP-PubMedBERT-base-uncased-abstract

2 https://huggingface.co/dmis-lab/biobert-large-cased-v1.1

3 https://dl.fbaipublicfiles.com/biolm/RoBERTa-large-PM-M3-Voc-hf.tar.gz

4 https://huggingface.co/sultan/BioM-ELECTRA-Large-Discriminator

5 https://huggingface.co/sultan/BioM-ALBERT-xxlarge

